# Structures of the Human SPAK and OSR1 Conserved C-Terminal (CCT) Domains

**DOI:** 10.1101/2021.08.19.456998

**Authors:** Karen T. Elvers, Magdalena Lipka-Lloyd, Rebecca C. Trueman, Benjamin D. Bax, Youcef Mehellou

**Affiliations:** Cardiff School of Pharmacy and Pharmaceutical Sciences, Cardiff University, Cardiff, CF10 3NB, U.K.; Medicines Discovery Institute, Cardiff University, Cardiff, CF10 3AT, U.K.; School of Life Sciences University of Nottingham Nottingham, NG7 2UH, U.K.

**Keywords:** SPAK, OSR1, Kinase, Crystal structure, Inhibitor

## Abstract

STE20/SPS1-related proline/alanine-rich kinase (SPAK) and Oxidative Stress Responsive 1 (OSR1) kinase are two serine/threonine protein kinase that regulate the function of ion co-transporters through phosphorylation. The highly conserved C-terminal (CCT) domains of SPAK and OSR1 bind to RFx[V/I] peptide sequences from their upstream With No Lysine Kinases (WNKs), facilitating their activation via phosphorylation. Thus, the inhibition of SPAK and OSR1 binding, via their CCT domains, to WNK kinases is a plausible strategy for inhibiting SPAK and OSR1 kinases. To facilitate structure-guided drug design of such inhibitors, we expressed and purified human SPAK and OSR1 CCT domains and solved their crystal structures. We also employed a biophysical strategy and determined the affinity of SPAK and OSR1 CCT domains to an 18-mer peptide derived from WNK4. Together, the crystal structures and affinity data reported herein provide a robust platform to facilitate the design of CCT domain specific small molecule inhibitors of SPAK-activation by WNK kinases, potentially leading to new improved treatments for hypertension and ischemic stroke.

## Introduction

The WNK-SPAK/OSR1 signalling pathway is a key regulator of ion homeostasis in humans.^[3]^ WNK kinases get activated under osmotic stress leading to the phosphorylation, and consequently, the activation of SPAK and OSR1 protein kinases **(Figure 1)**.^[4]^ SPAK and OSR1 kinases bind their upstream WNK kinases via their highly conserved *C*-terminal (CCT) domains, which bind to RFx[V/I] tetrapeptide sequences that are present in the four human WNK isoforms **(Figure 1)**. After binding and phosphorylation by WNK kinases, active SPAK and OSR1 kinases bind the scaffolding Mouse only protein 25 (MO25),^[5]^ of which there are two human isoforms α and β,^[6]^ and as a complex they phosphorylate a number of sodium, potassium and chloride ion co-transporters such as Na-K-2Cl co-transporters 1 and 2 (NKCC1 and 2), Na-Cl co-transporter and K-Cl co-transporter (KCC) **(Figure 1)**.^[3]^ Notably, the binding of SPAK and OSR1 kinases to these ion co-transporters is also mediated by the binding of their CCT domains to RFx[V/I] tetrapeptide motifs on these ion co-transporters akin to the binding of SPAK and OSR1 to WNK kinases **(Figure 1)**. Intriguingly, WNK kinases themselves have CCT-like (CCTL) domains that are have been reported to bind an GRFKV peptide,^[7]^ and the structures of CCTL domains from several WNK kinases have been solved (https://www.thesgc.org/tep/wnk3) and the literature relating to this has recently been reviewed.^[8]^

**Figure 1.**
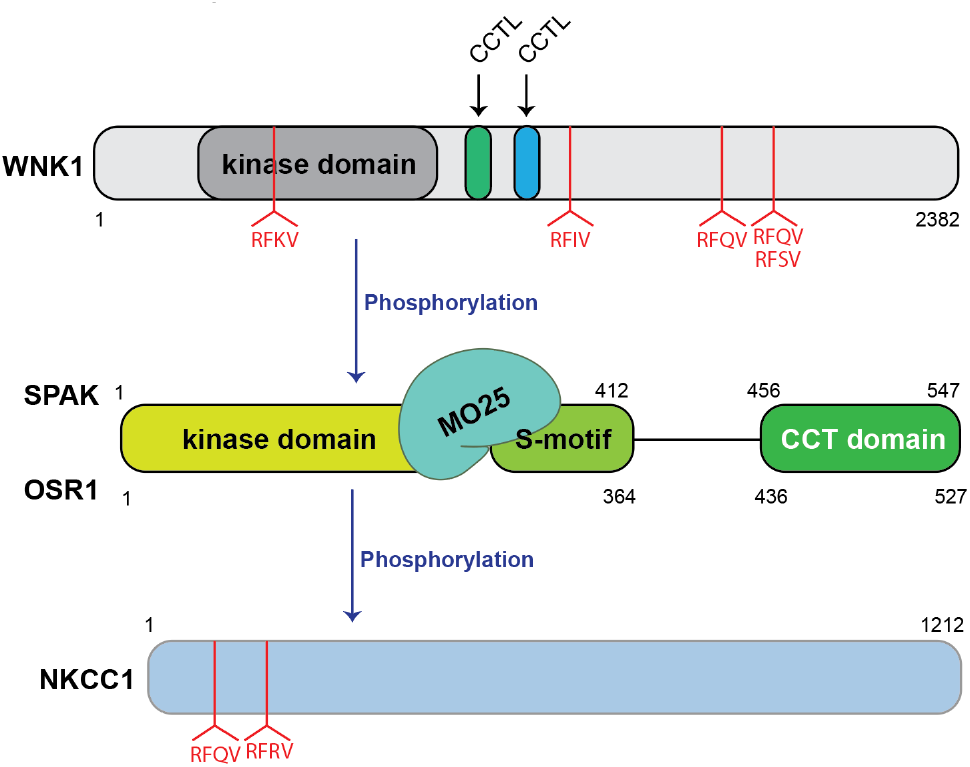
A schematic overview summary of the WNK-SPAK/OSR1-signalling pathway. The four human WNK kinase isoforms and the downstream ion co-transporter substrates of SPAK and OSR 1 kinases contain RFx[V/I] tetrapeptide motifs that mediate the binding to SPAK and OSR1 CCT domains, but for simplicity only WNK1 and NKCC1 are shown in this figure.

To date, there have been a series of reports in the literature documenting mutations of proteins that make up this signalling pathway resulting in various human diseases. For instance, genetic mutations in the WNK kinases have been found to lead to familial hyperkalemic hypertension; known as Gordon’s Syndrome.^[9]^ Additionally, mutations in two E3 ubiquitin ligases, Cullin 3 (Cul3) and Kelch-Like Protein 3 (KLHL3), which regulate the total protein levels of WNK kinases,^[10]^ have also been found to lead to familial hyperkalemic hypertension.^[11]^ Although most of the studies on the involvement of the WNK-SPAK/OSR1 signalling axis in human diseases have focused on hypertension, there is growing *in vivo* evidence of the involvement of this signalling pathway in ischemic strokes^[12]^ and breast cancer^[13]^.

A series of genetically modified WNK, SPAK and OSR1 animal models have shown that inhibition of this signalling pathway leads to a reduction in blood pressure,^[14]^ which offers beneficial outcomes in managing ischemic strokes^[15]^ and treating breast cancer^[13]^. Although this highlighted the therapeutic potential of inhibiting WNK-SPAK/OSR1 signalling, there have not been any small molecule inhibitors of this signalling cascade that progressed to clinical studies. Admittedly, there have been various inhibitors of WNK,^[16]^ SPAK and OSR1 kinases,^[15, 17-19]^ but these had limitations that hindered their further progress. Encouraged by the therapeutic potential of WNK-SPAK/OSR1 signalling inhibitors, this paper focuses on the interaction of SPAK and OSR1 CCT domains with the upstream WNK kinases, which has been shown *in vivo* to be critical for the function of this signalling cascade.^[20]^ In order to facilitate the discovery of small molecule binders of SPAK and OSR1 CCT domains, which inhibit their activation by the upstream WNK kinases, we solved crystal structure of both the human SPAK and OSR1 CCT domains.

## Results and Discussion

### Crystal structures of the human SPAK and OSR1 CCT domains

To obtain the first crystal structure of human SPAK CCT and a OSR1 CCT structure, we first designed and cloned new N-terminally His-tagged SPAK and OSR1 CCT constructs comprising amino acids 441-545 for SPAK and amino acids 423-527 for OSR1 (see Experimental Section for amino acid sequences). These proteins were then expressed in *E*.*coli*, purified and the His-tag was removed by protease digestion using standard methods (Supporting Figures S1 and S2, and also see Experimental). Crystal screens of the purified untagged proteins were set up. SPAK crystals were grown in 30 mM magnesium chloride, 30 mM calcium chloride, 50 mM sodium HEPES, 50 mM MOPS pH 7.5, 20% v/v PEG 500 MME and 10% w/v PEG 20000 at 6 °C, while OSR1 crystals were grown using 30 mM magnesium chloride, 30 mM calcium chloride, 50 mM sodium HEPES, 50 mM MOPS pH 7.5, 25% v/v MPD; 25% PEG 1000 and 25% w/v PEG 3350 at 6 °C. X-ray diffraction of these two crystals gave high resolution structures; a 1.73Å X-ray crystal structure of apo human SPAK CCT (PDB 7o86) and a 1.62Å apo structure of human OSR1 (PDB 7okw; Supporting Table S1 and **Figures 2A** and **2C**).

**Figure 2.**
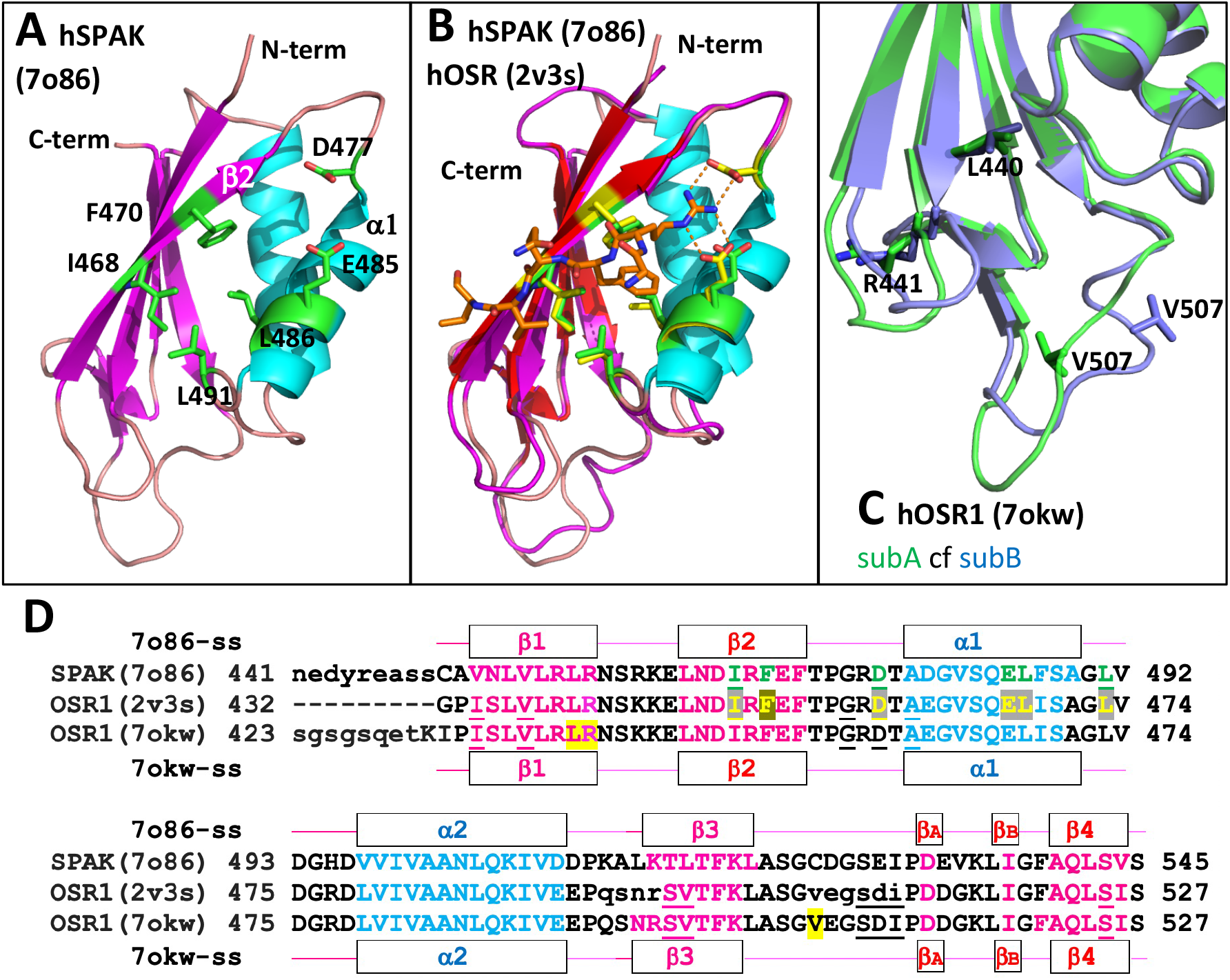
hSPAK and hOSR1 CCT crystal structures. **A**. 1.73Å structure of hSPAK1 CCT domain (PDB 7o86; this study) is shown as a cartoon. Six residues shown to be involved in peptide binding in OSR1 are shown as sticks with green carbons. **B**. The 1.70Å structure (PDB code 2v3s Villa *et al*.) of OSR1 with a peptide (GRFQVT; orange carbons –sticks) is shown superposed on **A**. The six residues I450, F452, D459, E467, L468 and L473 are shown as sticks with yellow carbons. **C**. A comparison of the ‘secondary pocket region’ between the two subunits in the 1.62Å hOSR1 crystal structure (PDB 7okw, this study). Side-chains of L440, R441 and V507 are shown in stick. **D**. Protein sequence alignment of the hSPAK CCT (7o86) and hOSR1 (2v3s and 7okw) CCT domains. The sequences are 69.5% identical. Secondary structural elements are shown above and below the amino-acid sequences. Lower case residues at the N-termini of the hSPAK (7o86) and hOSR1 (7okw) sequences indicate residues not observed in crystal structures. Lower case residues on the hOSR1 (2v3s) sequence are 494-497 and 507-512 which ‘*were disordered in the partial refined apo structure*’.^[1]^ Underlined residues were disordered in the NMR structure of hOSR1^[2]^.

Analysis of these two structures showed that, as expected, the SPAK CCT crystal structure was highly similar to that of the CCT domain of human OSR1 published by Villa *et al*.^[1]^ They published a 1.70Å OSR1 CCT domain (PDB 2v3s) in complex with a 6-mer GRFQVT-peptide corresponding to residues G1003 to T1008 from the human WNK4 (**Figure 2B**). The ‘primary’ pocket where the 6-mer peptide binds to OSR1 CCT (2v3s) is highly conserved in the SPAK (7o86) structure (and in the apo OSR1/7okw structure).

An unusual and apparently rigid part of the SPAK/OSR1 CCT domain structures contain a short β-bridge, containing two main-chain hydrogen bonds between Asp 532/514 and Ile 537/519 (these two residues are labelled as β-strands; βA and βB in **Figure 2D**, see also Supporting Figure S3). The side-chain of D532(SPAK) /D514(OSR1) also accepts hydrogen bonds from main-chain NHs of V534(SPAK)/G516(OSR1) and K535/K517; the structure between 532-537 SPAK (514-519 OSR) appears quite rigid.

In most OSR1/SPAK subunit structures the side-chain of R459 (SPAK)/R441 (OSR1), the last residue in the β1-strand, donates hydrogen bonds to two main-chain carbonyls of K535/517 and L536/518 (**Figure 2C**, Supporting Figure S3B). While R441 from the B subunit in the 1.62Å hOSR1 structure makes this ‘usual’ interaction (with main chain carbonyls of K517 and L518), in the A subunit an interaction is only observed between the side-chain of R441 and the carbonyl of K517. This move correlates with a change in the size of the ‘secondary pocket’,^[17]^ in the A subunit of the 1.62Å hOSR1 structure (**Figure 2C**).

In terms of the OSR1 structure, Villa *et al*.^[1]^ described an apo OSR1 CCT domain crystal structure, but this was not deposited in the PDB database due to relatively high R-factors (R=0.29, Rfree=0.35). Our deposited hOSR1 structure (7okw) and that of hOSR1 reported by Villa *et al*. (2v3s) are quite well defined in the 507-512 region (**Figure 2D**), but have differing structures in the secondary pocket region (**Figure 2C**). Notably, our 1.62Å OSR1 crystal was grown in the presence of a compound we identified as a possible inhibitor of SPAK and OSR1 binding to WNK kinases (unpublished data). The secondary pocket in subunit A in our structure (**Figure 2C**) has a different structure, which is stabilised by crystal contacts (our 7okw structure has two subunits in the asymmetric unit, called subunit A and subunit B; subunit B has a ‘normal’ secondary pocket structure). One possibility is that 507-512 of hSOR1 could form a hinge region allowing a C-terminal strand exchange.^[21]^

Although the SPAK CCT domain structure we have refined and deposited with the PDB (7o86) has reasonable R-factors (R=0.196, Rfree=0.240), the structure has high temperature factors (Supporting Figure S4B) in loops in similar positions to the OSR1 ‘disorderd loops’. Moreover, it is possible to model a ‘domain swapped’ dimer into the SPAK CCT structure (**Supporting Figure S4B**). One possibility could be that the rigid structure centred on the βA and βB strands (**Figure 2**) could be used as part of an allosteric mechanism to define different shapes of the ‘secondary pocket’.

### Comparison of the SPAK CCT domain with CCTL domains from WNK kinases

In 2013, the NMR solution structure of an autoinhibitory CCTL domain from WNK1, which associated with a GRFKV peptide, was reported.^[7]^ Although this domain had the conserved primary pocket peptide binding site observed in the SPAK and OSR1 CCT domain, the domain lacks the last two strands observed in OSR1 and SPAK CCT domains (**Figure 3**). The SGC have now solved crystal structures of number of CCTL domains from WNK kinases (https://www.thesgc.org/tep/wnk3), which all have the basic same fold as that observed by Moon *et al*^[7]^. One of these, a well refined 1.1Å structure WNK2 CCTL1domain (PDB 6elm) is compared with the SPAK CCT domain structure in **Figure 3**. Note that β-strands 3 and 4 from the 1.73Å SPAK structure are not observed in the WNK2 CCTL1, but a strand in the same position as β4 comes at the N-terminus of the domain (we call this strand β0) see **Figure 2**. The primary pocket is conserved between CCT and CCTL domains, but the secondary pocket is not conserved. Differences in the secondary pocket between the CCT domains of SPAK/OSR1 and the CCTL domains within WNK kinases may be important in the activation of SPAK/OSR1 by WNKs. Interestingly although the 1.1Å structure of the WNK2 CCTL1 domain has excellent R-factors (R = 0.138, Rfree = 0.168), a 1.7Å structure of the same WNK2 CCTL1 with an 18mer RFXV-motif peptide (PDB 6FBK) does not have such good R-factors (R = 0.284, Rfree = 0.334) (https://www.thesgc.org/tep/wnk3).

**Figure 3.**
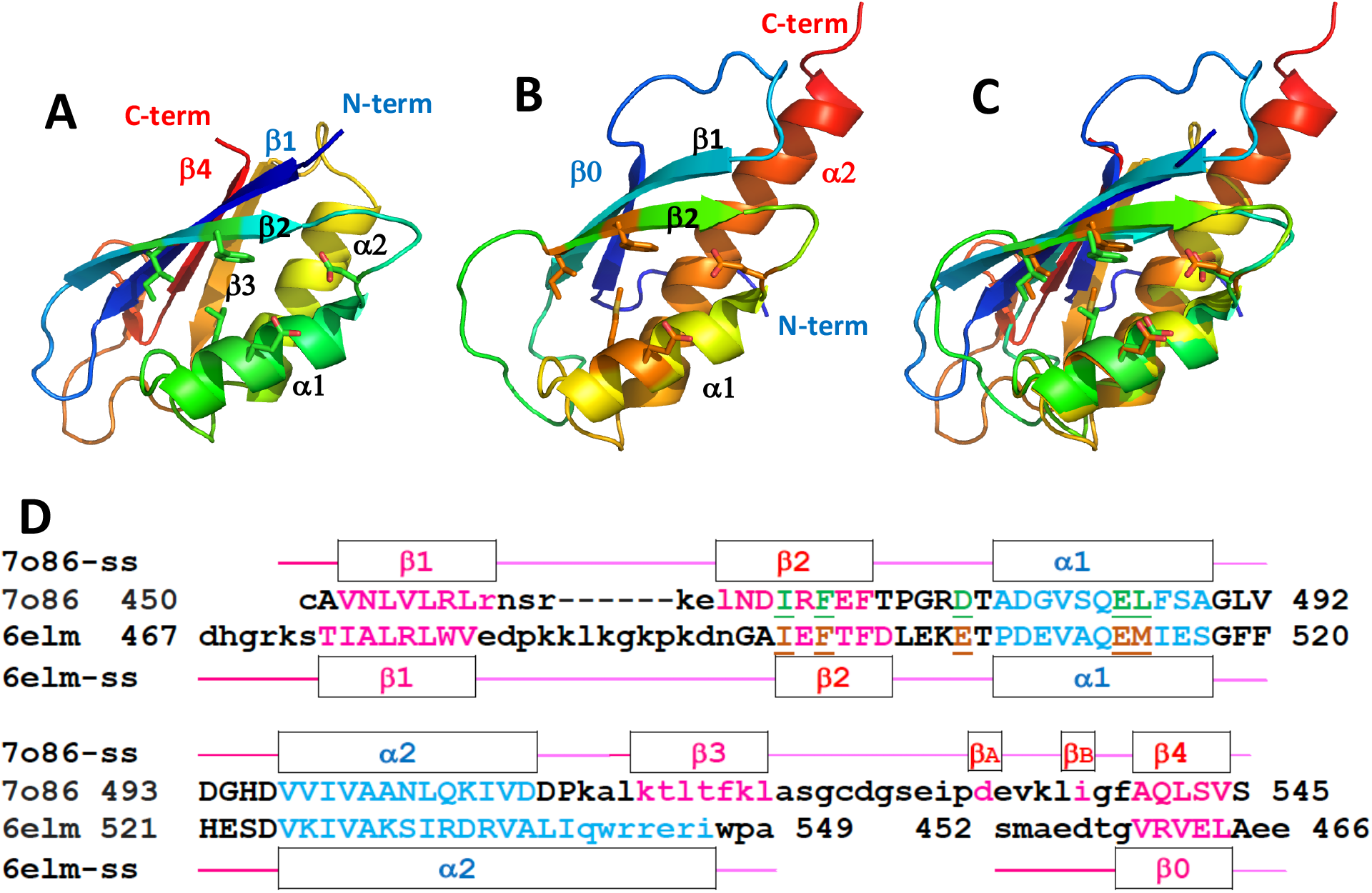
Comparison of the SPAK-CCT structure with the CCTL1 domain from WNK2. **A**. SPAK-CCT structure (pdb code 7o86) is shown as cartoon with colour ramp from blue (N-term – Cys 450) to red (C-term – Ser 545). The four strands in the β-sheet and the two α-helices are labelled as shown on sequence alignment. **B**. Wnk2 CCTL1 structure (pdb code 6elm) is shown as cartoon with colour ramp from blue (N-term – Ser 452) to red (C-term – Ala 549). The three strands in the β-sheet and the two α-helices are labelled as shown on sequence alignment. Note that the β0 strand comes before the β1 strand. **C** comparison of A and B. **D** a structurally based sequence alignment. Uppercase residues superpose – lowercase residues are in different positions in the two structures. Five residues, involved in forming the peptide binding pocket are underlined and coloured. These residues are shown as side-chain sticks in panels A and B.

### Analysis of binding properties of SPAK and OSR1 CCT domains to WNK kinases

As mentioned above, SPAK and OSR1 CCT domains interact with the upstream WNK kinases and the downstream ion co-transporters by binding to RFx[V/I] containing sequences. A number of studies investigated the binding affinities of peptides derived from WNK4 to SPAK and OSR1 CCT domains using a range of biophysical assays namely Surface Plasmon Resonance (SPR)^[22]^ and Fluorescence Polarisation (FP)^[17, 20]^. The binding affinity ranged from one study to another depending on the assay and conditions used (Supporting Table S2). Below, new biophysical assays that employ the Creoptix WAVE system (www.creoptix.com) are described. These assays have the advantage of offering insights into the kinetics of untagged SPAK and OSR1 CCT domains binding to an 18-mer RFQV peptide derived from WNK4. Notably, this 18-mer RFQV peptide was the same peptide used in an SPR assay by Vitari *et al*.^*[22]*^.

In the first instance, we embarked on measuring the binding affinity of the 18-mer RFQV peptide to SPAK and OSR1 CCT domains by labelling the proteins with biotin and immobilising them on a chip while the 18-mer peptide was the analyte, and the binding was evaluated using waveRAPID and traditional multicycle kinetics (**Figure 4**). The results showed that using a 1:1 kinetic model and global fitting, the K_D_ for the 18-mer peptide at 5 µM was 1.1 µM and 2.5 µM for SPAK and OSR1, respectively (**Figure 4A** and **4C**). The K_D_ value was essentially the same in waveRAPID when the 18-mer peptide, SPAK and OSR1 were used at different concentrations (Supporting Figures S5 and S6 and Supporting Table S3). Also, the K_D_ for the 18-mer was 1.16 and 2.5 µM for SPAK and OSR1 respectively with traditional multicycle kinetics (**Figure 4B** and **4D**). These K_D_ values are comparable to those reported previously (Supporting Table S2)^[17, 19, 20, 23]^ using SPR and fluorescence polarisation although most of these studies employed GST-tagged SPAK and OSR1 CCT domains, which will be dimeric. This may account for some of the differences observed in values in the literature. In order to verify that the measured binding affinity is a true reflection of the binding of the 18-mer peptide to SPAK and OSR1 CCT domains, we subsequently measured the binding affinity of the 18-mer peptide to SPAK L491A and OSR1 L473A. The results showed that there was no binding between the WNK-derived 18-mer to SPAK L491A and OSR1 L473A (Supporting Figures S7 and S8), and this is in agreement with the previous data obtained for measuring the binding affinity of these SPAK and OSR1 mutants to the RFQV 18-mer peptide (Supporting Table S2) ^[20, 22]^.

**Figure 4.**
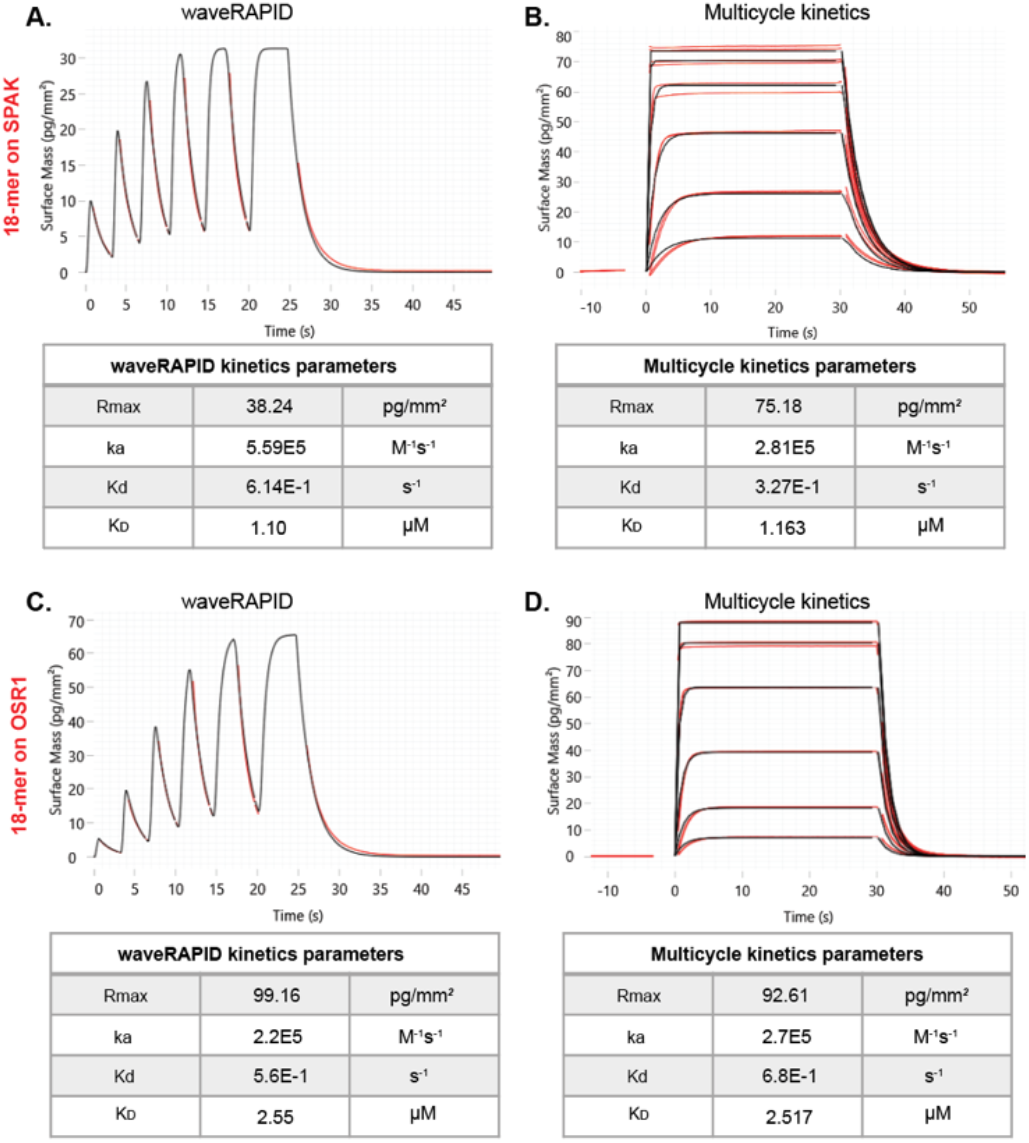
Comparison of multicycle kinetics with waveRAPID kinetics. Binding was measured for the 18-mer peptide SEEGKPQLVGRFQVTSSK (analyte) to immobilized biotin-SPAK or biotin-OSR generated with either waveRAPID (**A** and **C**, left panel; 18-mer at 5 µM) or with traditional multicycle kinetics (**B** and **D** right panel; 18-mer at 50 µM 3 fold dilutions). The double-referenced response data (red) are fitted with a one-to-one binding (black lines) with a suitable model in waveControl (model traditional fit). Table summaries of kinetic parameters are shown: Rmax, ka, association rate constant; kd, dissociation rate constant; and K_D_ dissociation constant.

Subsequently, we attempted measuring the binding affinity of the WNK-derived 18-mer RFQV peptide to SPAK and OSR1 CCT domains using the reverse situation where the *N*-terminally biotinylated 18-mer RFQV peptide was immobilised to the streptavidin surface and SPAK and OSR1 CCT domains were the mobile analytes. In this case, the binding affinity of OSR1 to the peptide resulted in K_D_ 2.1 and 6.6 µM using waveRAPID and multicycle methods respectively (**Figure 5B** and **5C**). For SPAK, there was clearly a more pronounced slow-off rate and/or non-specific binding component. In waveRAPID, this can be seen where the dissociation does not return quickly to baseline (Figure 5A). A K_D_ of 1.2 µM was determined using a heterogeneous model in which the heterogeneous ligand model supposes two types of differently interacting binding and is able to separate the two (**Figure 5A**). In multicycle kinetics, the incomplete dissociation of the slow binding component prevents the software fitting a model to the data (data not shown). In this scenario, waveRAPID achieves good data because it has a much lower contact time, which does not allow the slow component time to build up on the surface.

**Figure 5.**
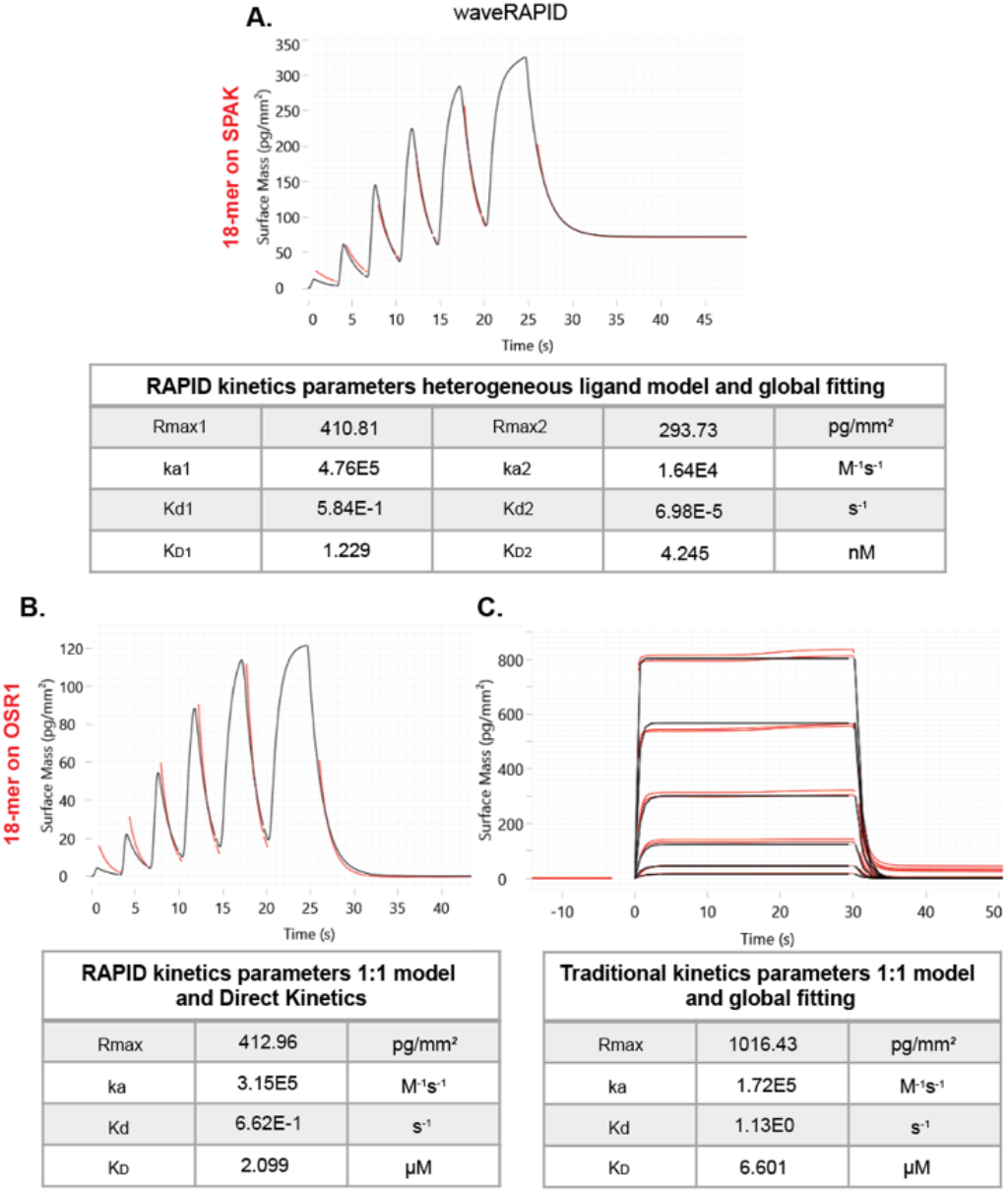
Comparison of traditional kinetic data and estimates with waveRAPID. Binding was measured for cleaved SPAK and OSR (analyte) to captured biotin-C_6_-SEEGKPQLVGRFQVTSSK generated with either waveRAPID (**A** for SPAK and **B** for OSR1) or with traditional multi-cycle kinetics (**C** for OSR1). The double-referenced response data (red) are fitted with a binding model (black lines). Here the binding model and fitting are indicated on the Figure. Table summaries of kinetic parameters are shown: Rmax 1 and 2, ka, association rate constant; kd, dissociation rate constant; and K_D_ dissociation constant.

Collectively, either of the two strategies used for measuring the binding affinities of the RFQV 18-mer peptide to SPAK and OSR1 CCT domains gives values that are comparable to previously reported K_D_ values. This validates the Creoptix WAVE method as a viable strategy for measuring the binding affinity of the RFQV 18-mer peptide to SPAK and OSR1 CCT domains and highlights the potential of using this assay for screening inhibitors of SPAK and OSR1 CCT domains binding to WNK kinases.

## Conclusion

In order to facilitate the discovery of SPAK and OSR1 kinase inhibitors, we herein reported the crystal structures of SPAK and OSR1 CCT domains. These domains are believed to be critical in the binding of SPAK and OSR1 kinases to their upstream WNK kinases, a process that leads to SPAK and OSR1 activation via phosphorylation by WNK kinases. After SPAK and OSR1 are activated by phosphorylation by WNK kinases, conformational changes are presumed to allow SPAK/OSR1 to then interact and phosphorylate downstream ion channels using the same type of RFx[V/I] CCT domain interaction.

The crystal structures of SPAK and OSR1 CCT domains will be powerful resources in the *in silico* design of new SPAK and OSR1 inhibitors. Additionally, the Creoptix WAVE assay reported in this work is also a powerful technology for screening small molecules that bind SPAK and OSR1 CCT domains, and in deciphering the fascinating regulatory interactions of the proteins with RFx[V/I] motifs from both upstream WNK kinases and downstream ion channels. Collectively, this work provides new resources and tools that will facilitate the discovery of new SPAK and OSR1 inhibitors that have the potential to ultimately improve the lifestyle of patients suffering from hypertension and who are at high risk or have had ischemic strokes.

## Experimental Section

### Construct design

Human SPAK and OSR1 CCT domain constructs were designed with: (i) an N-terminal His tag, (ii) an Avi-Tag (*GLNDIFEAQKIEWHE*) a unique 15 amino acid peptide tag for targeted enzymatic conjugation of a single biotin on the lysine using biotin ligase and (iii) a TEV-cleavage site (**ENLYFQG**). The DNAs for these were synthesised by GenScript (in pUC57). The amino acid sequence of the expressed uncleaved hSPAK construct (residues 441-545 underlined) is: MHHHHHHSS*GLNDIFEAQKIEWHE*SS**ENLYFQG**<u>NEDYREASSCAVNL VLRLRNSRKELNDIRFEFTPGRDTADGVSQELFSAGLVDGHDVVIVAANLQKIVDDPKALKTLTFKLASGCDGSEIPDEVKLIGFAQLSVS</u>. After cleavage with TEV protease the protein hSPAK-CCT contains a single additional glycine residue from the tag at the N-terminus (G*<u>NEDY</u>*..). The amino acid sequence of the human OSR1 (423-527 underlined) CCT is: HMHHHHHHSS*GLNDIFEAQKIEWHE*SS**ENLYFQG**<u>SGSGSQETKIPISL VLRLRNSKKELNDIRFEFTPGRDTAEGVSQELISAGLVDGRDLVIVAANLQKIVEEPQSNRSVTFKLASGVEGSDIPDDGKLIGFAQLSIS</u>. A further two constructs were synthesised by GenScript as above but with point mutations leucine to alanine (in bold and underlined) at amino acid L473 in OSR1 and L491in SPAK.

The amino acid sequence for uncleaved mutated hSPAK construct is: MHHHHHHSS*GLNDIFEAQKIEWHE*SS**ENLYFQG**<u>NEDYREASSCAVNLVLRLRNSRKELNDIRFEFTPGRDTADGVSQELFSAG</u>**<u>A</u>**<u>VDGHDVVIVAA NLQKIVDDPKALKTLTFKLASGCDGSEIPDEVKLIGFAQLSVS</u> and uncleaved mutated hOSR1 is: HMHHHHHHSS*GLNDIFEAQKIEWHE*SSENLYFQG<u>SGSGSQETKIPISLV LRLRNSKKELNDIRFEFTPGRDTAEGVSQELISAG</u>**<u>A</u>**<u>VDGRDLVIVAANL QKIVEEPQSNRSVTFKLASGVEGSDIPDDGKLIGFAQLSIS</u>

### Cloning and Expression

hSPAK-CCT, hOSR1-CCT, hSPAK L491A and hOSR1 L473A constructs were subcloned into a pET24a(+) expression vector using standard molecular methods, using OneShot TOP10 Chemically Competent *E. coli* (Invitrogen) and appropriate antibiotics. DNA sequences were confirmed by sequencing (Eurofins). For protein expression, plasmids were transformed into BL21-CodonPlus (DE3)-RIL competent *E. coli* (Agilent Technologies). Cultures was grown at 37°C to an OD600 0.6-0.8 in LB media with kanamycin (50µg/ml) and chloramphenicol (35 µg/ml). Expression was induced by the addition of 0.125 mM isopropyl β-D-1-thiogalactopyranoside (IPTG; Melford Laboratories Ltd), and the culture grown overnight at 20°C.

### Purification of hSPAK-CCT and hOSR-CCT

Uncleaved proteins (hSPAK-CCT, hOSR-CCT, L491A and L473A) were purified from supernatant of lysed cells, in two steps (at 4°C). 1. Affinity capture on a 5 ml HisTrap FF column in wash buffer [50 mM Tris.HCl pH 7.8, 300 mM NaCl, 5% glycerol and 5 mM imidazole], and eluted with a linear imidazole gradient [50 mM Tris.HCl pH 7.8, 300 mM NaCl, 5% glycerol and 250 mM imidazole]. 2. Size exclusion chromatography (SEC) on a Superdex 16/600 75 pg with SEC buffer [20 mM Tris.HCl pH 7.8, 50 mM NaCl, 1 mM DTT] (see Supporting Figures S1 and S2). Proteins were concentrated using MWCO 3K PES centrifugal concentrators (Pall Corporation) at 4,700xg at 10°C before loading on the SEC column. Uncleaved protein eluted as a single peak. Fractions were pooled, concentrated and subsequently quantified by Bradford assay. SDS-PAGE (16% Tris-Glycine; Invitrogen) was used for analysis at each stage.

### Modifications post purification

#### i. SPAK and OSR1 for crystallisation

SPAK-CCT was cleaved at a Q/G site with Tobacco Etch Virus (TEV) Protease, to remove the N-terminal 6xHis (underlined) and AviTag (bold) (M<u>HHHHHH</u>SS**GLNDIFEAQKIEWHE**SS*ENLYFQ*/G). The cleaved tag was removed by incubation with 0.5 ml Super Cobalt NTA Agarose Affinity Resin (Generon) and the flow through containing SPAK was loaded on and run on a Superdex 16/600 75pg SEC column pre-equilibrated with SEC buffer [20 mM Tris.HCl pH 7.8, 50 mM NaCl, 1 mM DTT]. Fractions were pooled and concentrated using Cerocon (SWISSCI) 0.5ml concentrator at 3 kDa MWCO. They were quantified by Bradford assay.

#### ii. SPAK and OSR1 as analytes for surface-based biophysical measurement of binding kinetics

Both proteins were cleaved with TEV as above, but the final SEC was run using HBS EP buffer [10mM HEPES pH7.4, 150 mM NaCl, 3 mM EDTA, 0.005% Tween20].

#### iii. Biotinylated SPAK and OSR1 and alanine mutations as ligands for binding kinetics

Enzymatic biotinylation with *E. coli* biotin ligase (BirA) was carried out to specifically biotinylate the lysine residue in the AviTag sequence GLNDIFEAQ**K**IEWHE. Briefly, 750 µg of purified uncleaved hSPAK-CCT or hOSR-CCT were mixed with 50 µl of SuperMix (containing ATP and D-biotin, Avidity, LLC) and 5 μg of BirA (Avidity, LLC), to a final volume of 500 μL buffer [20 mM Tris.HCl pH 7.8, 50 mM NaCl, 1 mM DTT]. The reaction was incubated overnight at 4 °C with mixing. The free biotin and BirA ligase were removed from the final biotinylated proteins by size exclusion in HBS EP buffer. Final fractions containing biotinylated protein were concentrated and quantified by Bradford assay and termed biotin-SPAK and biotin-OSR.

### Crystallisation, data collection and structure determination of CCTs

Cleaved hSPAK-CCT (1.4 mg/ml) was crystallised using a Mosquito liquid handling robot (TTP Labtech) in a SWISSCI 3 lens sitting drop plates, using a drop ratio of 150 nls protein (in 20 mM Tris.HCl pH 7.8, 50 mM NaCl, 1 mM DTT) and 50 nls well. The SPAK crystal structure reported in this paper was from a crystal grown against well of Morpheus A5 (30 mM magnesium chloride, 30 mM calcium chloride, 50 mM sodium HEPES, 50 mM MOPS pH 7.5, 20% v/v PEG 500 MME, 10% w/v PEG 20000) at 6 °C. The crystal used for data collection was picked in a loop and plunged into liquid nitrogen. Data were collected on beam-line i03 at Diamond Light Source. The 1.73Å dataset (supplementary Table 1) was autoprocessed with DIALS ^[24]^ run via the xia2 autoprocessing pipeline ^[25]^. The dataset contains some 3600, 0.1 degree images (360 degrees of data); examination of merging statistics by batch did not show any major variation with batch. The Rmerge in low resolution shells is around 6%.

The structure was solved by molecules replacement with phaser ^[26]^ using the OSR1 CCT (PDB code: 2v3s, residues 434-526 of subunit A – some side-chains trimmed). The structure was refined with Refmac5 ^[27]^ and phenix.refine ^[28]^ and rebuilt in coot ^[29]^. Coordinates have been deposited with the PDB (pdb code: 7o86).

Cleaved hOSR1-CCT (1.89 mg/ml) was crystallised using a Mosquito liquid handling robot (TTP Labtech) in a SWISSCI 3 lens sitting drop plates, using a drop ratio of 150nls protein (in 20 mM Tris.HCl pH 7.8, 50 mM NaCl, 1 mM DTT + compound 1 at 1 mM and 10%DMSO) and 50 nls well. The OSR1 crystal structure reported in this paper was grown against well of Morpheus A8 (30 mM magnesium chloride, 30 mm calcium chloride, 50 mM sodium HEPES, 50 mM MOPS pH 7.5, 25% v/v MPD; 25% PEG 1000; 25% w/v PEG 3350) at 6 °C.

The crystal used for data collection was picked in a loop and plunged into liquid nitrogen. Data were collected on beam-line i04 at Diamond Light Source. The 1.62Å dataset (Supporting Table S1) was autoprocessed with Autoproc/STARANISO.^[30]^ The diffraction limits of ellipsoid fitted to diffraction cut-off surface were 1.598, 1.763 and 1.768Å. The structure was solved by molecules replacement with phaser ^[26]^ using the OSR1 CCT (PDB code: 2v3s, residues 434-526 of subunit A - some side-chains trimmed). The structure was refined with Refmac5^[27]^ and phenix.refine^[28]^ and rebuilt in coot ^[29]^. Coordinates have been deposited with the PDB (pdb code: 7okw).

### Structure analysis

Secondary structures were calculated with DSSP^[31]^ as implemented on the web server https://swift.cmbi.umcn.nl/gv/dssp/.^[32]^ The only helices shown in Figures are α-helices. Buried area between different subunits in crystals were analysed with the program PISA^[33]^ at the EBI (https://www.ebi.ac.uk/pdbe/pisa/).

### Surface-based biophysical measurement of binding kinetics

Grated-Coupled Interferometry (GCI) experiments were performed on a Creoptix WAVE system (Creoptix, AG) using 4PCH STA WAVE sensor chips (polycarboxylate surface, streptavidin coated). Chips were conditioned with borate buffer (100 mM sodium borate pH9, 1 M NaCl; Xantec). Biotin-SPAK, biotin-OSR1, biotin-SPAK L491A and biotin-OSR1 L473A were directly immobilized onto the sensor chip by injection onto the surface in HBS EP [10 mM HEPES pH7.4, 150 mM NaCl, 3 mM EDTA, 0.005% Tween20] buffer at 10 µg/ml at a flowrate of 10 µl/min. The final surface density for the biotin-SPAK, biotin-OSR and alanine mutations was about 1000 pg/mm^2^. The biotinylated 18-mer peptide (Biotin-C_6_-SEEGKPQLVGRFQVTSSK) was purchased from (GL Biochem, Shanghai Ltd). The density of the biotin-peptide was controlled to about 400 pg/mm^2^. Kinetic analyses were performed at 25°C using, **(a)** waveRAPID (Repeated Analyte Pulses of Increasing Duration - multiple short pulses of the analyte at a single concentration but of increasing durations.^[34]^ The experiments were carried out at flow rate of 100 µl/min. Briefly, 122 µL of samples were applied to the immobilised surface and reference for 25 s total injection duration, followed by a 300 s dissociation with assay buffer and **(b)** traditional multicycle kinetics where the analyte is introduced at increasing concentrations with injection of uniform duration. The experiment was carried out at flow rate of 120 µl/min. 110 µL of the analytes were injected for 30 s, followed by a 300 s dissociation for 18mer peptide and OSR and 2000 s dissociation for SPAK with assay buffer.

The analyses were measured with two scenarios.

i. immobilised biotin-SPAK or biotin-OSR, or biotin-SPAK L491A or biotin-OSR1 L473A as ‘ligands’ with binding kinetics of unbiotinylated 18 mer peptide (SEEGKPQLVGRFQVTSSK, GL Biochem, Shanghai Ltd).
ii. the inverse set up, immobilised biotin 18-mer peptide as the ‘ligand’ and the binding kinetics of cleaved SPAK and cleaved OSR1 were analysed. The following concentrations of analytes were used.
  i. <u>Immobilised SPAK or OSR</u>: In waveRAPID; the 18-mer peptide concentration was 2, 5, 10 or 20 µM. In multicycle kinetics, the 18 mer was serially diluted 3-fold using assay buffer for a six point dose curve (50 µM, 17 µM, 5.5 µM, 1.8 µM, 0.6 µM, 0.2 µM).
  ii. <u>Immobilised 18 mer peptide</u>: In waveRAPID when biotinylated peptide was captured onto the chip the kinetics of cleaved OSR and SPAK at 2 µM. In multicycle kinetics cleaved OSR and SPAK were serially diluted 1:3 using assay buffer for a six point dose curve (25 µM, 8.3 µM, 2.8 µM, 925.9 nM, 308.6 nM, 102.9 nM).

Blank injections were used for double referencing and a DMSO calibration curve for bulk correction. Analysis and correction of the obtained data were performed using the Creoptix WAVE control software 4.2 (correction applied: X and Y offset; DMSO calibration; double referencing). One-to-one binding and heterogeneous models were used with a suitable fitting model.

## Acknowledgements

We thank Fabio Andres and Edward FitzGerald (Creoptix AG, Wädeswil, Switzerland) for their excellent support in the analysis of the Creoptix WAVE data and suggestions for additional experiments. The work was funded by a Medical Research Council grant (Ref. 518455) awarded to Y.M. and B.D.B.

## Entry for the Table of Contents

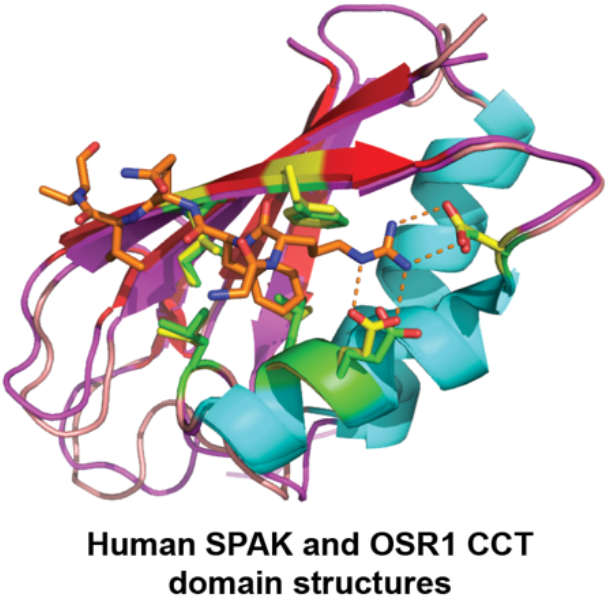

The inhibition of SPAK and OSR1 kinases binding to their upstream WNK kinases has been identified as a plausible strategy for inhibiting the WNK-activation of SPAK and OSR1 kinases. To facilitate the discovery of SPAK and OSR1 kinases inhibitors as potential therapeutics, we herein report the crystal structures of the highly conserved C-terminal domains of SPAK and OSR1, which mediate the binding to, and activation, by their upstream WNK kinases.

Institute and/or researcher Twitter usernames: @Youcef_Mehellou, @PharmacyCU, @CUMedicinesInst

## Notes

### Competing Interest Statement

The authors have declared no competing interest.

## References

[1] F. Villa, J. Goebel, F. H. Rafiqi, M. Deak, J. Thastrup, D. R. Alessi, D. M. van Aalten, EMBO Rep. 2007, 8, 839–845.

[2] M. A. AlAmri, M. Jeeves, Y. Mehellou, Biochem. Biophys. Res. Commun. 2019, 512, 338–343.

[3] a)D. R. Alessi, J. Zhang, A. Khanna, T. Hochdorfer, Y. Shang, K. T. Kahle, Sci. Signaling 2014, 7, re3; b)J. Hadchouel, D. H. Ellison, G. Gamba, Ann. Rev. Physiol. 2016, 78, 367–389.

[4] A. C. Vitari, M. Deak, N. A. Morrice, D. R. Alessi, Biochem. J.2005, 391, 17–24.

[5] B. M. Filippi, P. de los Heros, Y. Mehellou, I. Navratilova, R. Gourlay, M. Deak, L. Plater, R. Toth, E. Zeqiraj, D. R. Alessi, EMBO J. 2011, 30, 1730–1741.

[6] J. Boudeau, A. F. Baas, M. Deak, N. A. Morrice, A. Kieloch, M. Schutkowski, A. R. Prescott, H. C. Clevers, D. R. Alessi, EMBO J. 2003, 22, 5102–5114.

[7] T. M. Moon, F. Correa, L. N. Kinch, A. T. Piala, K. H. Gardner, E. J. Goldsmith, J. Mol. Biol. 2013, 425, 1245–1252.

[8] C. A. t. Taylor, M. H. Cobb, Molecular pharmacology 2021.

[9] F. H. Wilson, S. Disse-Nicodeme, K. A. Choate, K. Ishikawa, C. Nelson-Williams, I. Desitter, M. Gunel, D. V. Milford, G. W. Lipkin, J. M. Achard, M. P. Feely, B. Dussol, Y. Berland, R. J. Unwin, H. Mayan, D. B. Simon, Z. Farfel, X. Jeunemaitre, R. P. Lifton, Science 2001, 293, 1107–1112.

[10] a)A. Ohta, F. R. Schumacher, Y. Mehellou, C. Johnson, A. Knebel, T. J. Macartney, N. T. Wood, D. R. Alessi, T. Kurz, Biochem. J. 2013, 451, 111–122; b)M. Wakabayashi, T. Mori, K. Isobe, E. Sohara, K. Susa, Y. Araki, M. Chiga, E. Kikuchi, N. Nomura, Y. Mori, H. Matsuo, T. Murata, S. Nomura, T. Asano, H. Kawaguchi, S. Nonoyama, T. Rai, S. Sasaki, S. Uchida, Cell Rep. 2013, 3, 858–868; c)S. Shibata, J. Zhang, J. Puthumana, K. L. Stone, R. P. Lifton, Proc. Nat. Acad. Sci. USA 2013, 110, 7838–7843.

[11] a)H. Louis-Dit-Picard, J. Barc, D. Trujillano, S. Miserey-Lenkei, N. Bouatia-Naji, O. Pylypenko, G. Beaurain, A. Bonnefond, O. Sand, C. Simian, E. Vidal-Petiot, C. Soukaseum, C. Mandet, F. Broux, O. Chabre, M. Delahousse, V. Esnault, B. Fiquet, P. Houillier, C. I. Bagnis, J. Koenig, M. Konrad, P. Landais, C. Mourani, P. Niaudet, V. Probst, C. Thauvin, R. J. Unwin, S. D. Soroka, G. Ehret, S. Ossowski, M. Caulfield, P. Bruneval, X. Estivill, P. Froguel, J. Hadchouel, J. J. Schott, X. Jeunemaitre, Nature Genet. 2012, 44, 456–460; b)L. M. Boyden, M. Choi, K. A. Choate, C. J. Nelson-Williams, A. Farhi, H. R. Toka, I. R. Tikhonova, R. Bjornson, S. M. Mane, G. Colussi, M. Lebel, R. D. Gordon, B. A. Semmekrot, A. Poujol, M. J. Valimaki, M. E. De Ferrari, S. A. Sanjad, M. Gutkin, F. E. Karet, J. R. Tucci, J. R. Stockigt, K. M. Keppler-Noreuil, C. C. Porter, S. K. Anand, M. L. Whiteford, I. D. Davis, S. B. Dewar, A. Bettinelli, J. J. Fadrowski, C. W. Belsha, T. E. Hunley, R. D. Nelson, H. Trachtman, T. R. Cole, M. Pinsk, D. Bockenhauer, M. Shenoy, P. Vaidyanathan, J. W. Foreman, M. Rasoulpour, F. Thameem, H. Z. Al-Shahrouri, J. Radhakrishnan, A. G. Gharavi, B. Goilav, R. P. Lifton, Nature 2012, 482, 98–102.

[12] a)G. Begum, H. Yuan, K. T. Kahle, L. Li, S. Wang, Y. Shi, B. E. Shmukler, S. S. Yang, S. H. Lin, S. L. Alper, D. Sun, Stroke 2015, 46, 1956–1965; b)H. Zhao, R. Nepomuceno, X. Gao, L. M. Foley, S. Wang, G. Begum, W. Zhu, V. M. Pigott, L. M. Falgoust, K. T. Kahle, S. S. Yang, S. H. Lin, S. L. Alper, T. K. Hitchens, S. Hu, Z. Zhang, D. Sun, J. Cereb. Blood Flow Metab. 2017, 37, 550–563.

[13] Y. Li, L. Li, J. Qin, J. Wu, X. Dai, J. Xu, Oncogene 2021, 40(1), 68–84; Y. Li, J. Qin, J. Wu, X. Dai, J. Xu, Breast Cancer Res. Treat. 2020, 182, 35-46.

[14] F. H. Rafiqi, A. M. Zuber, M. Glover, C. Richardson, S. Fleming, S. Jovanovic, A. Jovanovic, K. M. O’Shaughnessy, D. R. Alessi, EMBO Mol. Med. 2010, 2, 63–75.

[15] J. Zhang, M. I. H. Bhuiyan, T. Zhang, J. K. Karimy, Z. Wu, V. M. Fiesler, J. Zhang, H. Huang, M. N. Hasan, A. E. Skrzypiec, M. Mucha, D. Duran, W. Huang, R. Pawlak, L. M. Foley, T. K. Hitchens, M. B. Minnigh, S. M. Poloyac, S. L. Alper, B. J. Molyneaux, A. J. Trevelyan, K. T. Kahle, D. Sun, X. Deng, Nature cCommun. 2020, 11, 78.

[16] a)K. Yamada, H. M. Park, D. F. Rigel, K. DiPetrillo, E. J. Whalen, A. Anisowicz, M. Beil, J. Berstler, C. E. Brocklehurst, D. A. Burdick, S. L. Caplan, M. P. Capparelli, G. Chen, W. Chen, B. Dale, L. Deng, F. Fu, N. Hamamatsu, K. Harasaki, T. Herr, P. Hoffmann, Q. Y. Hu, W. J. Huang, N. Idamakanti, H. Imase, Y. Iwaki, M. Jain, J. Jeyaseelan, M. Kato, V. K. Kaushik, D. Kohls, V. Kunjathoor, D. LaSala, J. Lee, J. Liu, Y. Luo, F. Ma, R. Mo, S. Mowbray, M. Mogi, F. Ossola, P. Pandey, S. J. Patel, S. Raghavan, B. Salem, Y. H. Shanado, G. M. Trakshel, G. Turner, H. Wakai, C. Wang, S. Weldon, J. B. Wielicki, X. Xie, L. Xu, Y. I. Yagi, K. Yasoshima, J. Yin, D. Yowe, J. H. Zhang, G. Zheng, L. Monovich, Nature Chem. Biol. 2016, 12, 896–898; b)K. Yamada, J. H. Zhang, X. Xie, J. Reinhardt, A. Q. Xie, D. LaSala, D. Kohls, D. Yowe, D. Burdick, H. Yoshisue, H. Wakai, I. Schmidt, J. Gunawan, K. Yasoshima, Q. K. Yue, M. Kato, M. Mogi, N. Idamakanti, N. Kreder, P. Drueckes, P. Pandey, T. Kawanami, W. Huang, Y. I. Yagi, Z. Deng, H. M. Park, ACS Chem. Biol. 2016, 11, 3338–3346; c)K. Yamada, J. Levell, T. Yoon, D. Kohls, D. Yowe, D. F. Rigel, H. Imase, J. Yuan, K. Yasoshima, K. DiPetrillo, L. Monovich, L. Xu, M. Zhu, M. Kato, M. Jain, N. Idamakanti, P. Taslimi, T. Kawanami, U. A. Argikar, V. Kunjathoor, X. Xie, Y. I. Yagi, Y. Iwaki, Z. Robinson, H. M. Park, J. Med. Chem. 2017, 60, 7099–7107.

[17] M. A. AlAmri, H. Kadri, L. J. Alderwick, N. S. Simpkins, Y. Mehellou, ChemMedChem 2017, 12, 639–645.

[18] E. Kikuchi, T. Mori, M. Zeniya, K. Isobe, M. Ishigami-Yuasa, S. Fujii, H. Kagechika, T. Ishihara, T. Mizushima, S. Sasaki, E. Sohara, T. Rai, S. Uchida, J. Am. Soc. Nephrol. 2015, 26, 1525–1536.

[19] T. Mori, E. Kikuchi, Y. Watanabe, S. Fujii, M. Ishigami-Yuasa, H. Kagechika, E. Sohara, T. Rai, S. Sasaki, S. Uchida, Biochem. J. 2013, 455, 339–345.

[20] J. Zhang, K. Siew, T. Macartney, K. M. O’Shaughnessy, D. R. Alessi, Human Mol. Genet. 2015, 24, 4545–4558.

[21] Y. Liu, D. Eisenberg, Protein Sci. 2002, 11, 1285–1299.

[22] A. C. Vitari, J. Thastrup, F. H. Rafiqi, M. Deak, N. A. Morrice, H. K. Karlsson, D. R. Alessi, Biochem. J. 2006, 397, 223–231.

[23] C. A. t. Taylor, S. W. An, S. G. Kankanamalage, S. Stippec, S. Earnest, A. T. Trivedi, J. Z. Yang, H. Mirzaei, C. L. Huang, M. H. Cobb, Proc. Nat. Acad. Sci. USA 2018, 115, 3840–3845.

[24] G. Winter, D. G. Waterman, J. M. Parkhurst, A. S. Brewster, R. J. Gildea, M. Gerstel, L. Fuentes-Montero, M. Vollmar, T. Michels-Clark, I. D. Young, N. K. Sauter, G. Evans, Acta Crystallogr. D Struct. Biol. 2018, 74, 85–97.

[25] G. Winter, C. M. Lobley, S. M. Prince, Acta Crystallogr. D Biol. Crystallogr. 2013, 69, 1260–1273.

[26] A. J. McCoy, Methods Mol. Biol. 2017, 1607, 421–453.

[27] O. Kovalevskiy, R. A. Nicholls, F. Long, A. Carlon, G. N. Murshudov, Acta Crystallogr. D Struct. Biol. 2018, 74, 215–227.

[28] P. D. Adams, P. V. Afonine, G. Bunkoczi, V. B. Chen, I. W. Davis, N. Echols, J. J. Headd, L. W. Hung, G. J. Kapral, R. W. Grosse-Kunstleve, A. J. McCoy, N. W. Moriarty, R. Oeffner, R. J. Read, D. C. Richardson, J. S. Richardson, T. C. Terwilliger, P. H. Zwart, Acta Crystallogr. D. Biol. Crystallogr. 2010, 66, 213–221.

[29] P. Emsley, B. Lohkamp, W. G. Scott, K. Cowtan, Acta Crystallogr. D. Biol. Crystallogr. 2010, 66, 486–501.

[30] C. Vonrhein, I. J. Tickle, C. Flensburg, P. Keller, W. Paciorek, A. Sharff, G. Bricogne, Acta Crystallogr. 2018, 74, a360–a360.

[31] W. Kabsch, C. Sander, Biopolymers 1983, 22, 2577–2637.

[32] W. G. Touw, C. Baakman, J. Black, T. A. te Beek, E. Krieger, R. P. Joosten, G. Vriend, Nucleic acids research 2015, 43(Database issue), D364–368.

[33] E. Krissinel, K. Henrick, J. Mol. Biol. 2007, 372, 774–797.

[34] Ö. Kartal, F. Andres, M. P. Lai, R. Nehme, K. Cottier, SLAS Discov. 2021, 24725552211013827.

