## Supporting Information for "Structures of the Human SPAK and OSR1 Conserved C-Terminal (CCT) Domains"

### CONTENTS

- **Supporting Table S1.** Data collection and refinement statistics.
- **Supporting Table S2.** Literature reporting KD or $IC_{50}$ values by different methods to the RFVQ motif of either WNK4 or WNK1.
- **Supporting Table S3.** Kinetic values for waveRAPID binding of the 18-mer peptide to SPAK and OSR1.
- **Supporting Figure S1.** SDS PAGE gels of purified of SPAK and OSR1.
- **Supporting Figure S2.** Electron density maps of the $\beta$-bridge between Asp532 (D532) and Ile537 (I537) in SPAK.
- **Supporting Figure S3.** A model of a strand exchanged dimer in the 1.73Å SPAK CCT domain structure.
- **Supporting Figure S4.** Size-exclusion chromatography elution profile of uncleaved SPAK-CCT with inset showing SDS-PAGE of peaks.
- **Supporting Figure S5.** Binding of 18-mer peptide to SPAK on the WAVE.
- **Supporting Figure S6.** Binding of 18-mer peptide to OSR1 on the WAVE.
- **Supporting Figure S7.** No binding of 18-mer peptide to SPAK L491A on the WAVE.
- **Supporting Figure S8.** No binding of 18-mer peptide to OSR1 L473A on the WAVE.

| PDB code: | 7o86 (SPAK-CCT) | 7okw (OSR1-CCT) |
|---|---|---|
| Diffraction source | IO3 – Diamond Light Source | IO4 – Diamond Light Source |
| Wavelength | 0.97628 | 0.97950 |
| Resolution range | 45.45 - 1.73 (1.792 - 1.73) | 55.7-1.62 (1.764-1.62) |
| Space group | P $2_1$ $2_1$ $2_1$ | C2 |
| Unit cell | 39.61 50.55 103.80 90 90 90 | 70.17 49.42 56.44 90 99.44 90 |
| Total reflections | 293450 (28311) | 132058 (6432) |
| Unique reflections | 22482 (2168) | 19062 (954) |
| Multiplicity | 13.1 (12.8) | 6.9 (6.7) |
| Completeness (%) | 99.63 (97.88) | 78.6 (17.9) ( spherical )<br>91.5 (49.2) ( ellipsoidal ) |
| Mean I/sigma(I) | 6.83 (0.43) | 10.0 (1.4) |
| Wilson B-factor | 25.83 | 21.04 |
| R-merge | 0.2553 (3.974) | 0.114 (1.246) |
| R-meas | 0.2657 (4.139) | 0.123 (1.352) |
| R-pim | 0.07308 (1.15) | 0.047 (0.519) |
| CC1/2 | 0.997 (0.281) | 0.999 (0.624) |
| Reflections used in refinement | 22400 (2165) | 19058 (360) |
| Reflections used for R-free | 1100 (106) | 930 (14) |
| R-work | 0.1965 (0.3791) | 0.1840 (0.3016) |
| R-free | 0.2400 (0.3820) | 0.2165 (0.2968) |
| CC(work) | 0.950 (0.621) | 0.967 (0.810) |
| CC(free) | 0.958 (0.517) | 0.913 |
| Number of non-hydrogen atoms | 1937 | 1733 |
| macromolecules | 1729 | 1532 |
| ligands | 4 | 21 |
| solvent | 204 | 180 |
| Protein residues | 191 | 194 |
| RMS(bonds) | 0.017 | 0.006 |
| RMS(angles) | 1.44 | 0.88 |
| Ramachandran favored (%) | 97.33 | 98.95 |
| Ramachandran allowed (%) | 2.14 | 1.05 |
| Ramachandran outliers (%) | 0.53 | 0.00 |
| Rotamer outliers (%) | 4.21 | 2.40 |
| Clashscore | 8.36 | 0.96 |
| Average B-factor | 34.40 | 25.71 |
| macromolecules | 33.58 | 24.55 |
| ligands | 51.69 | 46.03 |
| solvent | 41.04 | 33.24 |

Values in parentheses are for the outer shell. Note that the OSR1-CCT data were processed with STARANISO,[1] which processes data anisotropically.

**Supporting Table S1. Data collection and refinement statistics.**

| Reference | Construct | Method | KD (and $IC_{50}$ where indicated) |
|---|---|---|---|
| Vitari et al. 2006[2] | GST-OSR1 CCT (429-527)<br><br>GST-OSR1 L437A | Biacore; immobilised biotinylated RFQV peptide | 8 nM<br><br><br>No binding |
| Villa et al. 2007[3] | GSR-OSR1 (434-527)<br><br><br>GST-OSR1 L473A | Biacore; immobilised biotinylated RFQV peptide | Not cited. Fig 4A response units vs concentration<br><br><br>No binding |
| Zhang et al. 2015[4] | GST-SPAK (452-545)<br><br>GST-SPAK L491A | Fluorescence polarisation Lumino-Green-labelled WNK peptide | 0.6 µM<br><br>88.2 µM |
| Mori et al. 2013[5] | Rat GST-SPAK (452-553) | Fluorescent TAMRA labelled RFQV-WNK4<br>RFQV-WNK1<br>NCC-RFTI<br>AFQV-WNK4<br>Biacore; immobilised GST-SPAK with STOCK1S-50699 (PubChem-CID 5749625)<br>GST-SPAK with STOCK2S-26016 (PubChem-CID3135086) | 1.3 µM<br><br>1.3 µM<br>11.2 µM<br>No binding<br><br><br><br>32 µM<br><br><br><br>20 µM |
| Ishigami-Yuasa et al. 2017[6] | Rat GST-SPAK (452-553) | Fluorescent TAMRA RFQV-WNK4 and various substitutions in lead compound 1<br>Compound 10<br>Compound 13<br>Compound 20 | <br><br><br>15.4 µM $IC_{50}$<br>6.9 µM $IC_{50}$<br>2.6 µM $IC_{50}$<br>4.8 µM $IC_{50}$ |
| Kikuchi et al. 2015[7] | GST-SPAK [T233E] | Biacore 1S-14279 binding and ELISA $IC_{50}$<br><br><br>Closantel | KD not cited, $IC_{50}$ 0.26 µM<br><br>KD not cited, $IC_{50}$ 0.77 µM |
| AlAmri et al. 2017[8] | GST-OSR1 (433-end) | Fluorescence polarisation RFQV-WNK4 peptide | <br><br>2.11 µM |

| | | | |
|---|---|---|---|
| | GST-OSR1 L473A (433-end) | RFQV-WNK4 peptide | 13.17 µM |
| Taylor et al 2018[9] | His6-OSR CCT (433–527)<br><br>His6-SPAK CCT (449–545) | Fluorescence anisotrophy FAM probe chimera of WNK4 and WNK1 R-F-x-V peptide with competing WNK1 Probe vs WNK1 | 5.1 ± 0.4 µM<br><br>2.6 ± 0.2 µM |

**Supporting Table S2. Literature reporting $K_D$ or $IC_{50}$ values by different methods to the RFVQ motif of either WNK4 or WNK1.**

| Ligand | Analyte (conc) | Ka M-1s-1 | Kd s-1 | KD µM | Rmax pg/mm2 | Sqrt(chi2) pg/mm2 |
|---|---|---|---|---|---|---|
| SPAK | 18mer(2µM) | 475783.2 | 0.567316 | 1.19 | 43.62 | 0.17 |
| SPAK | 18mer(5µM) | 558691.7 | 0.613815 | 1.10 | 38.24 | 0.17 |
| SPAK | 18mer(10µM) | 552270.2 | 0.6317 | 1.14 | 34.74 | 0.18 |
| SPAK | 18mer(20µM) | 514495.5 | 0.705448 | 1.37 | 37.82 | 0.21 |
| OSR | 18mer(2uM) | 119380.5 | 0.515295 | 4.32 | 158.96 | 0.17 |
| OSR | 18mer(5uM) | 219859.4 | 0.560134 | 2.55 | 99.16 | 0.25 |
| OSR | 18mer(10uM) | 245014.4 | 0.571064 | 2.33 | 85.41 | 0.27 |
| OSR | 18mer(20uM) | 239972.6 | 0.618801 | 2.58 | 89.68 | 0.29 |

**Supporting Table S3. Kinetic values for waveRAPID binding of the 18-mer peptide to SPAK and OSR1.** Kinetic values for waveRAPID binding of the 18-mer peptide SEEGKPQLVGRFQVTSSK (analyte) at 2, 5, 10 and 20 µM, to immobilized biotin-OSR1 and biotin-SPAK on the Creoptix WAVE (see supplementary Figures 5 and 6 for raw data). Ka = association rate, kd = dissociation rate, $K_D$ = dissociation rate constant, Rmax = the maximum signal generated by an interaction between a ligand – analyte in pg/mm$^2$.

**A**

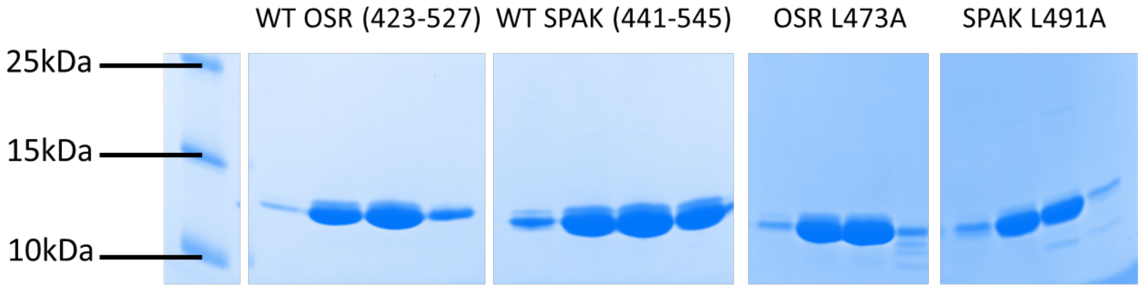


**B**

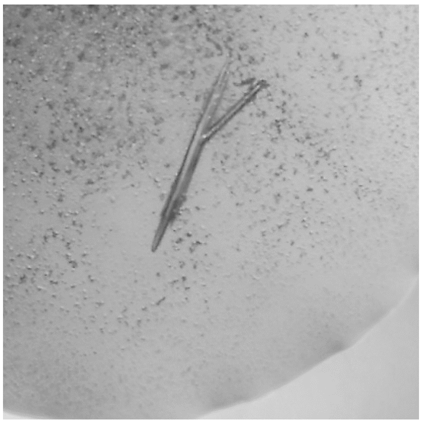

**Supporting Figure S1. SDS PAGE gels of purified of SPAK and OSR1. A**. SDS PAGE using 16% Tris Tricine gels of wild type human SPAK CCT, OSR1 CCT, SPAK [L491A] and OSR [L473A] after final size exclusion. The fractions for each were pooled and concentrated. Concentrated protein was either cleaved with TEV protease for crystallisation or uncleaved protein was biotinylated with BirA ligase for binding kinetics. Marker molecular weights are indicated in kD. **B**. Crystal of human SPAK from Morpheus screen A5 grown at 6°C.

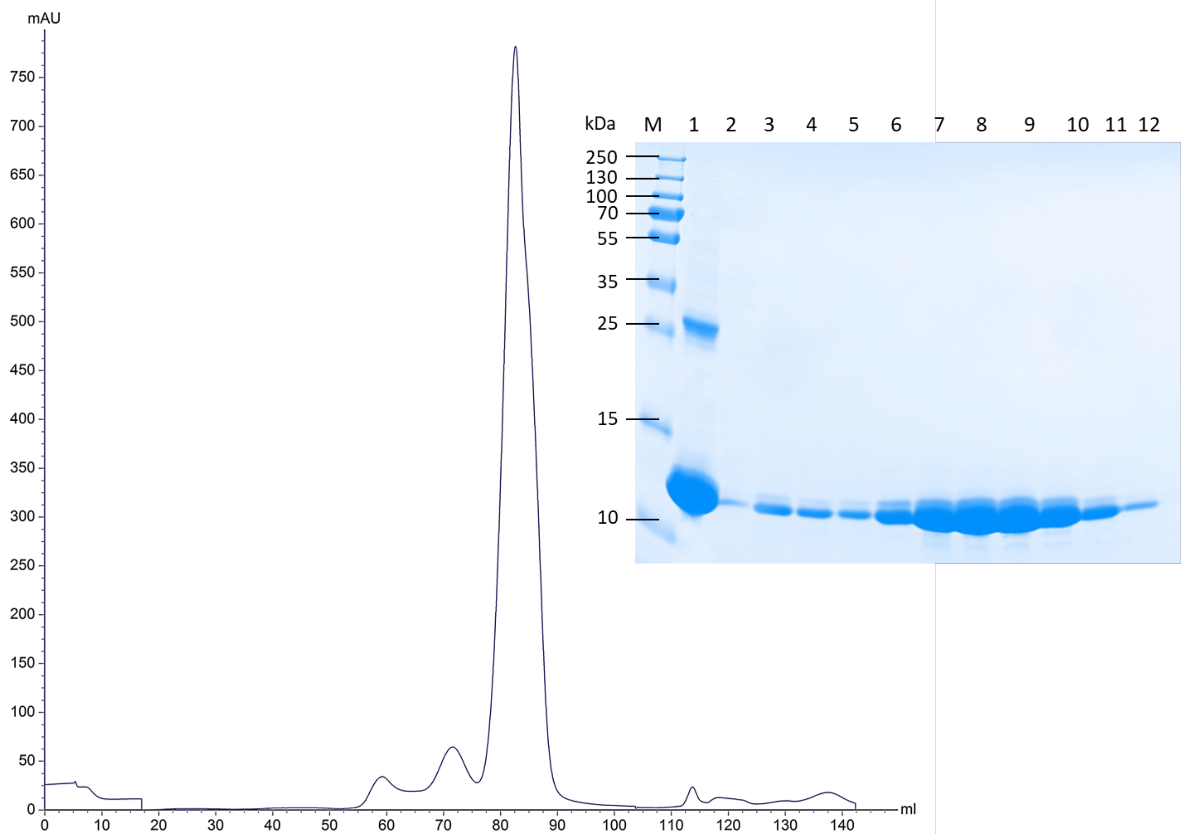


**Supporting Figure S2. Size-exclusion chromatography elution profile of uncleaved SPAK CCT with inset showing a 16% SDS-PAGE analysis of the peaks**. 'M' is protein markers. Lane 1 is the concentrated protein after HisTrap purification; Lanes 2 and 3 a fraction from each of the smaller peaks preceding the main peak. Lanes 4-12 protein contained in the fractions over the major peak. These were pooled and concentrated for cleavage for crystallography.

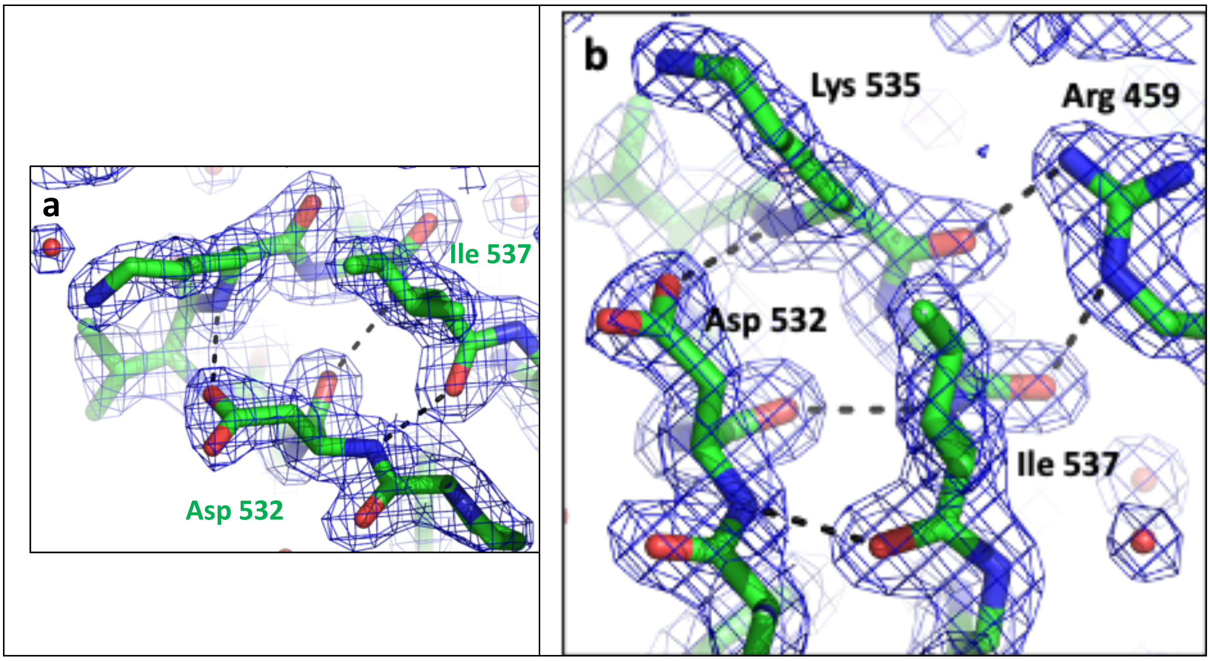


**Supporting Figure S3. Electron density maps of the b-bridge between Asp 532 (D532) and Ile 537 (I537) in SPAK**. **A**. The final 2fo-fc map (contoured at 1.4sigma) around Asp 532 (D532) and Ile 537 (I537) from the 1.73Å hSPAK structure. Dotted black lines are hydrogen bonds from involving Asp 532 (D532). **B**. An alternative view at 1.5sigma, showing hydrogen bonds to the side-chain of Arg 459 (R459), the C-terminal residue of the first β-strand (β1).

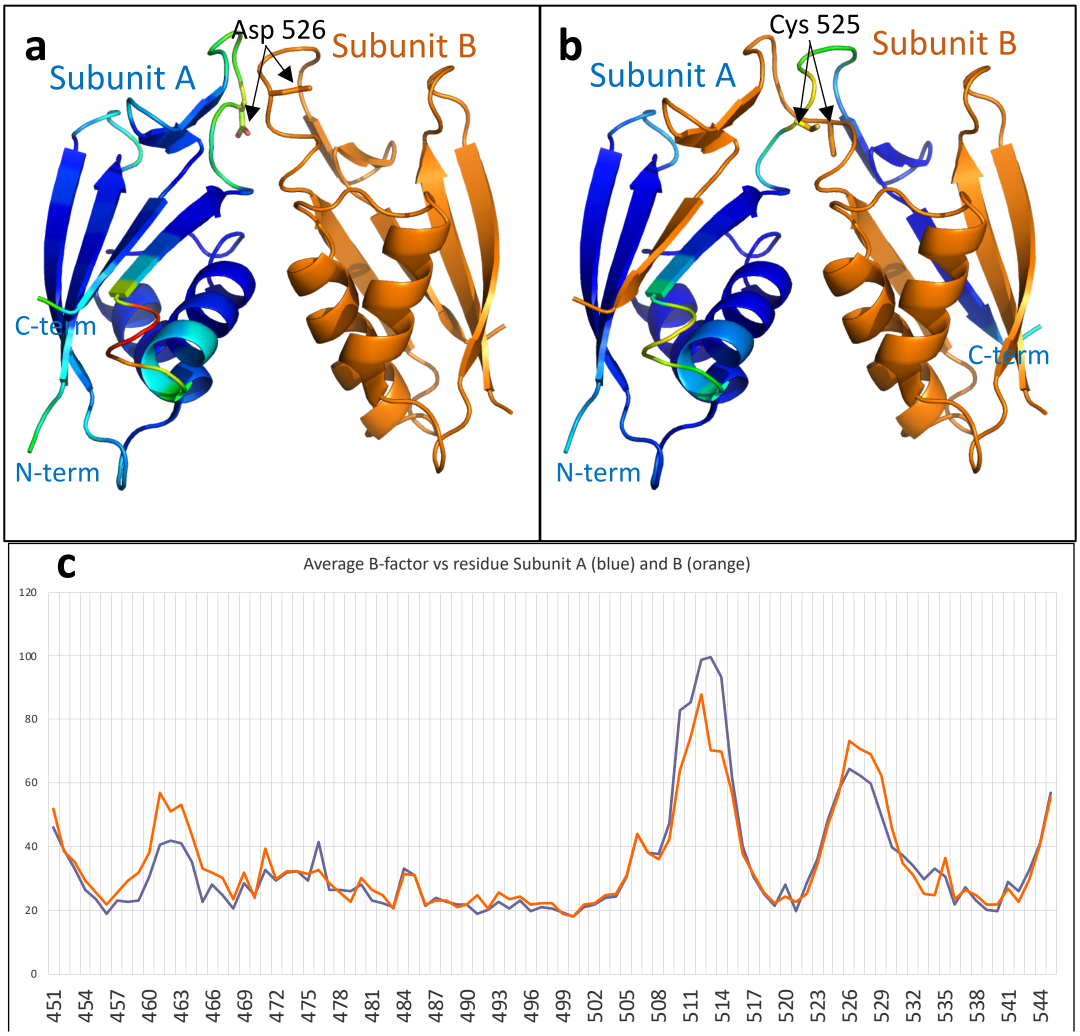


**Supporting Figure S4. A model of a strand exchanged dimer in the 1.73Å SPAK CCT domain structure. A**. 1.73Å structure of hSPAK1 (pdb code: 7o86) is shown as a cartoon. Subunit A is coloured by temperature factor (blue – cold, yellow – warm, red – hot). Subunit B is coloured orange. Side-chains of residue Asp 526 (D526) are shown on each subunit. **B**. An alternatively refined 'domain swapped' dimer of the same structure. Side-chains of Cys 525 (C525) are shown on each subunit. **C.** Average B factor versus residue number plot with subunit A shown by blue line and subunit B by orange line.

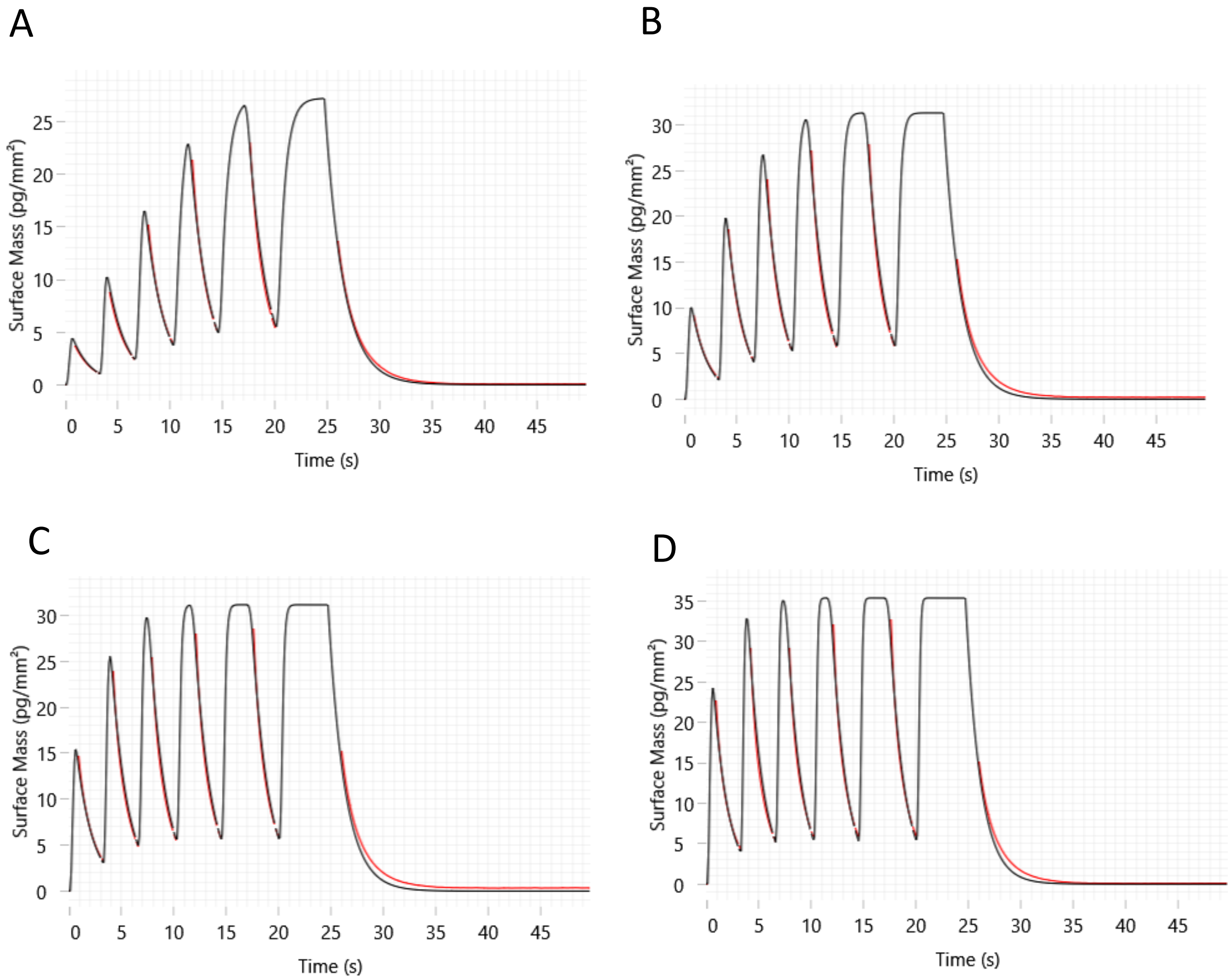


**Supporting Figure S5. Binding of 18-mer peptide to SPAK on the WAVE.** WaveRAPID kinetics. Binding was measured for the 18-mer peptide SEEGKPQLVGRFQVTSSK (analyte) at (**A)** 2 µM, (**B**) 5 µM, (**C**) 10 µM and (**D**) 20 µM to immobilized biotin-SPAK. The double-referenced response data (red) are fitted with a one-to-one binding model (black lines) in waveControl (see Supplementary Table 3 for derived kinetics values).

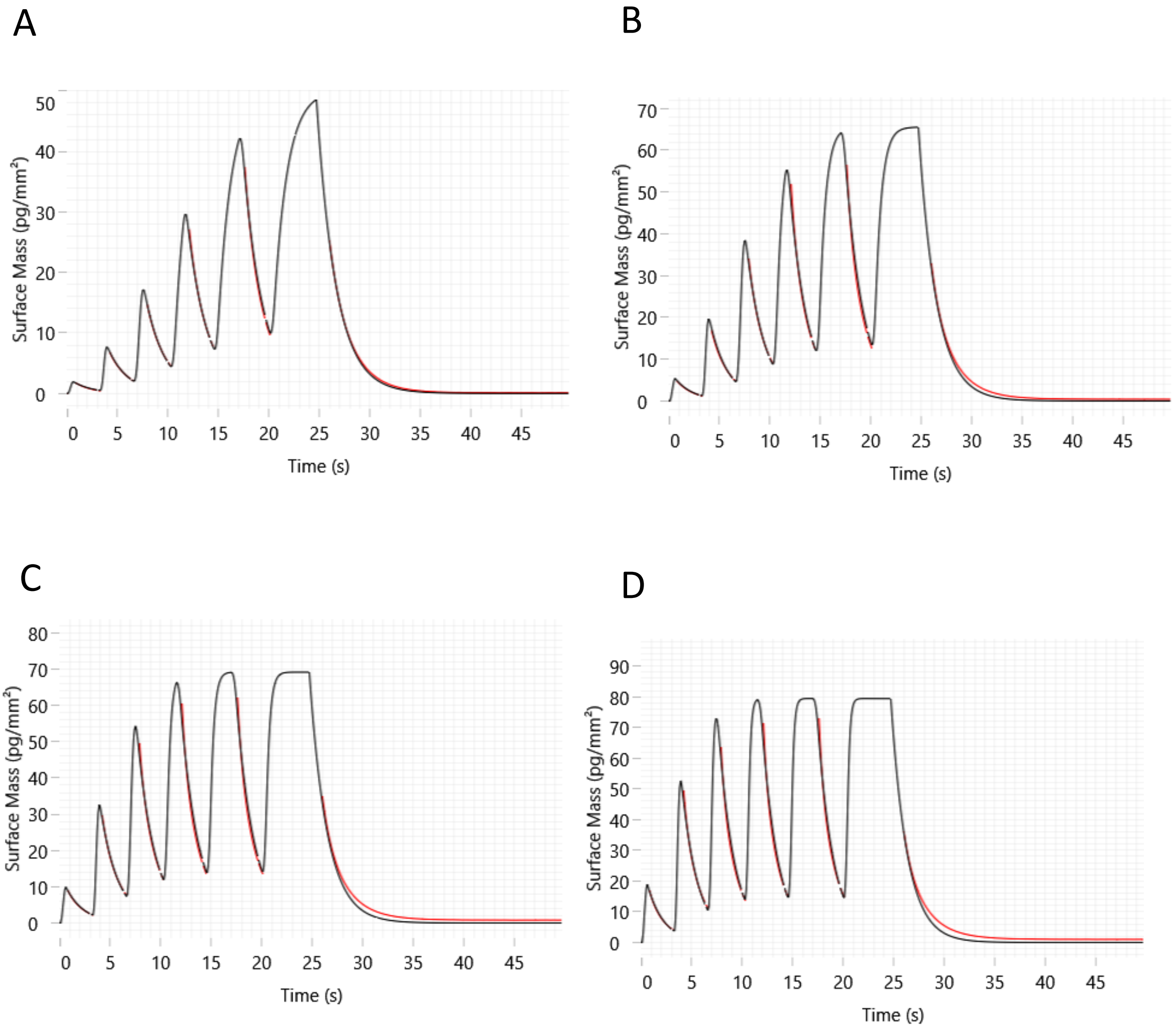


**Supporting Figure S6. Binding of 18-mer peptide to OSR1 on the WAVE.** WaveRAPID kinetics. Binding was measured for the 18-mer peptide SEEGKPQLVGRFQVTSSK (analyte) at (**A**) 2 µM, (**B**) 5 µM, (**C**) 10 µM and (**D**) 20 µM to immobilized biotin-OSR1. The double-referenced response data (red) were fitted with a one-to-one binding (black lines) with a suitable model in waveControl (see Supplementary Table 3 for derived kinetics values).

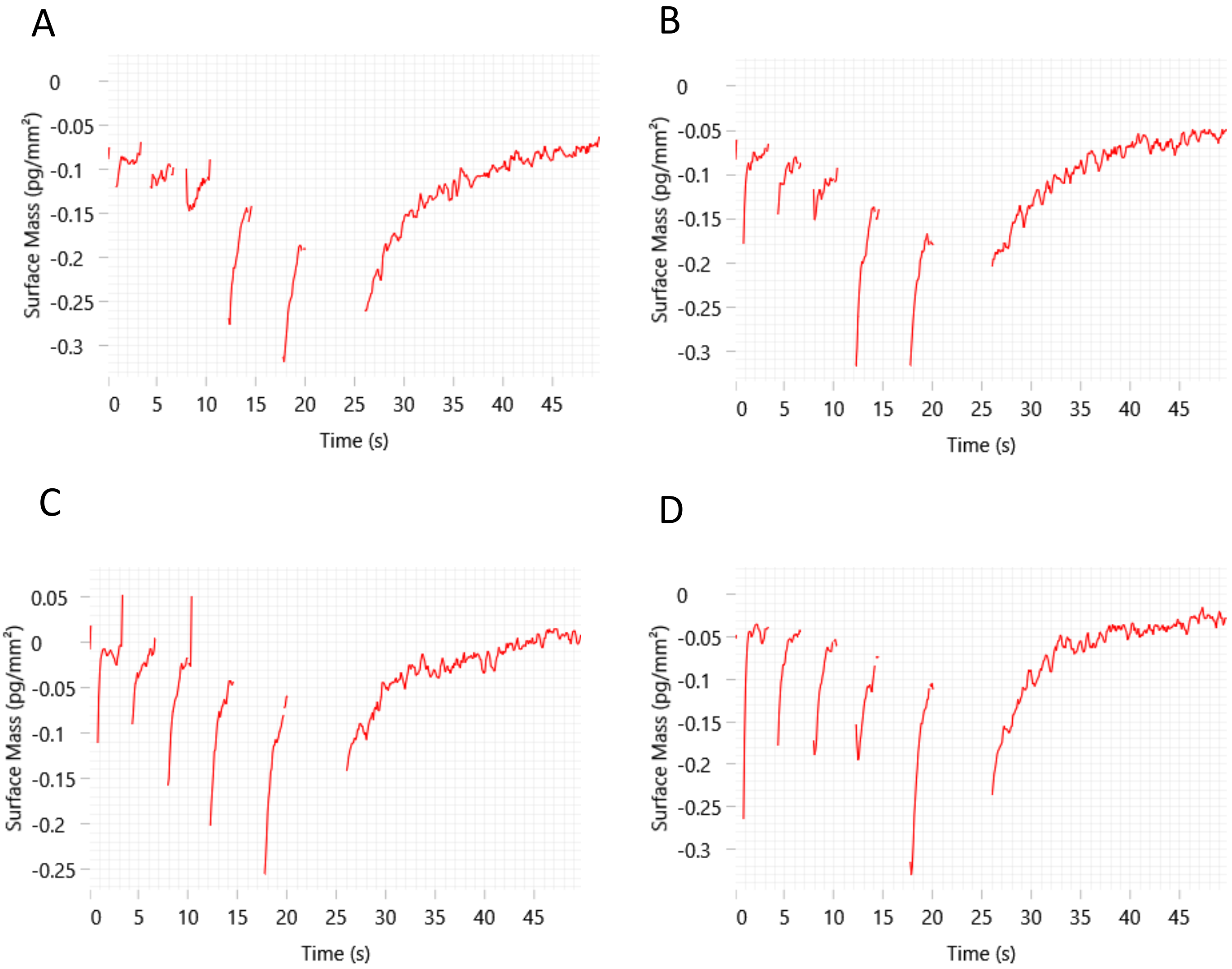


**Supporting Figure S7. No binding of 18-mer peptide to SPAK L491A on the WAVE.** WaveRAPID kinetics. Binding was measured for the 18-mer peptide SEEGKPQLVGRFQVTSSK (analyte) at (**A**) 2 µM, (**B**) 5 µM, (**C**) 10 µM and (**D**) 20 µM to immobilized biotin-SPAK L491A. The double-referenced response data (red) could noto be fitted with a suitable model in waveControl. Binding is not observed indicated by the high association and dissociation errors and no Rmax. Compare with supplementary Figure 5 - for binding of peptide to wild-type SPAK.

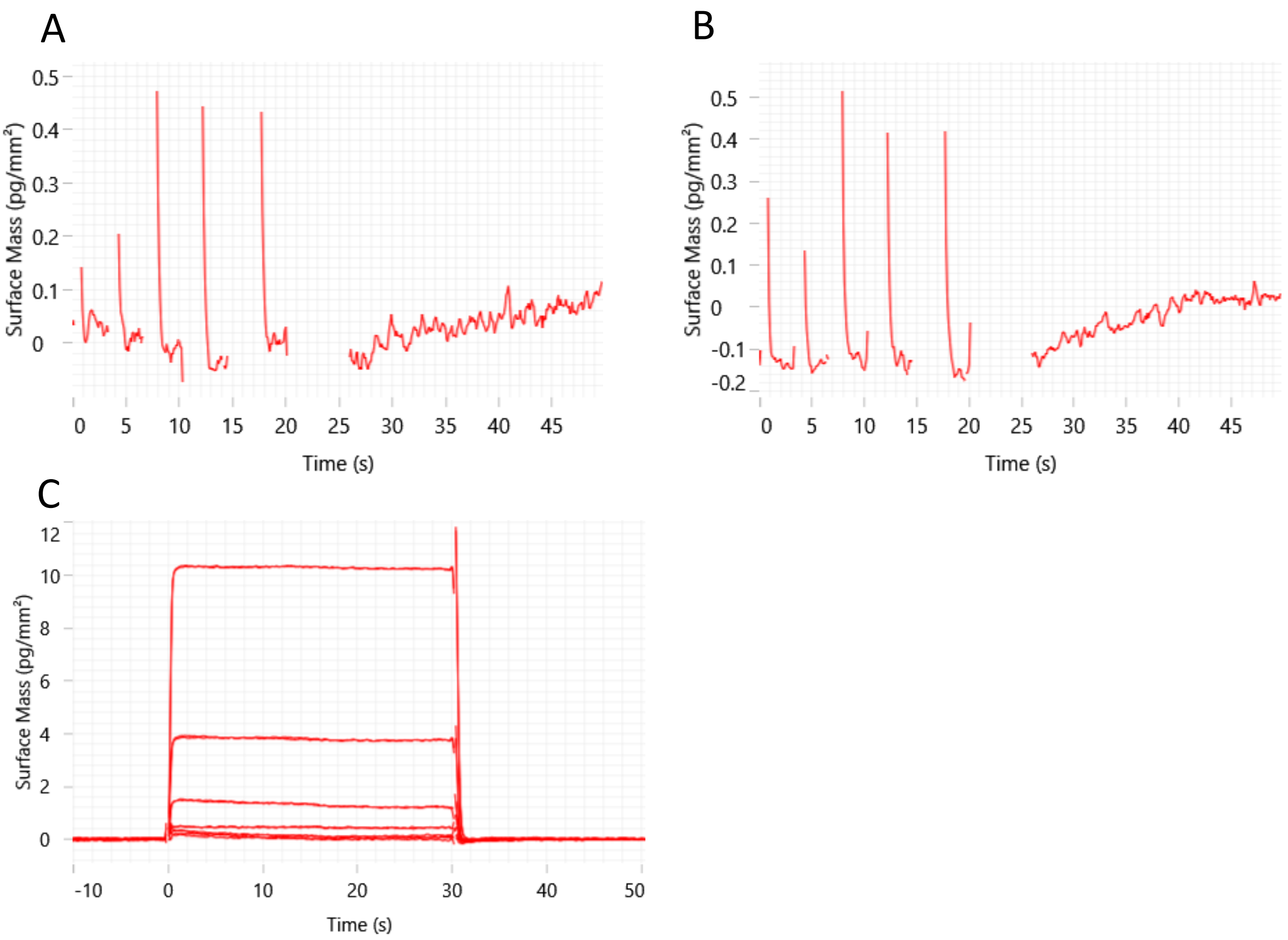


**Supporting Figure S8. No binding of 18-mer peptide to OSR1 L473A on the WAVE.** Raw data for binding of the 18-mer peptide SEEGKPQLVGRFQVTSSK (analyte) at (**A**) 10µM, (**B**) 20µM, to immobilized biotin-OSR1 L473A. (**C**) Multicylce kinetics for (50 µM, 17 µM, 5.5 µM, 1.8 µM, 0.6 µM, 0.2 µM) 18-mer peptide binding to OSR1 L473A. The double-referenced response data (red) could not be fitted with a one-to-one binding with a suitable model in waveControl. Binding is not observed indicated by the high association and dissociation errors and no Rmax. Compare with Supplementary Figure 6 for binding of 18-mer peptide to wild-type OSR1.

**References**


[1] C. Vonrhein, I. J. Tickle, C. Flensburg, P. Keller, W. Paciorek, A. Sharff, G. Bricogne, *Acta Crystallogr.* **2018**, *74*, a360-a360.

[2] A. C. Vitari, J. Thastrup, F. H. Rafiqi, M. Deak, N. A. Morrice, H. K. Karlsson, D. R. Alessi, *Biochem. J.* **2006**, *397*, 223-231.

[3] F. Villa, J. Goebel, F. H. Rafiqi, M. Deak, J. Thastrup, D. R. Alessi, D. M. van Aalten, *EMBO Rep.* **2007**, *8*, 839-845.

[4] J. Zhang, K. Siew, T. Macartney, K. M. O'Shaughnessy, D. R. Alessi, *Human Mol. Genet.* **2015**, *24*, 4545-4558.

[5] T. Mori, E. Kikuchi, Y. Watanabe, S. Fujii, M. Ishigami-Yuasa, H. Kagechika, E. Sohara, T. Rai, S. Sasaki, S. Uchida, *Biochem. J.* **2013**, *455*, 339-345.

[6] M. Ishigami-Yuasa, Y. Watanabe, T. Mori, H. Masuno, S. Fujii, E. Kikuchi, S. Uchida, H. Kagechika, *Bioorg. Med. Chem.* **2017**, *25*, 3845-3852.

[7] E. Kikuchi, T. Mori, M. Zeniya, K. Isobe, M. Ishigami-Yuasa, S. Fujii, H. Kagechika, T. Ishihara, T. Mizushima, S. Sasaki, E. Sohara, T. Rai, S. Uchida, *J. Am. Soc. Nephrol.* **2015**, *26*, 1525-1536.

[8] M. A. AlAmri, H. Kadri, L. J. Alderwick, N. S. Simpkins, Y. Mehellou, *ChemMedChem* **2017**, *12*, 639-645.

[9] C. A. t. Taylor, S. W. An, S. G. Kankanamalage, S. Stippec, S. Earnest, A. T. Trivedi, J. Z. Yang, H. Mirzaei, C. L. Huang, M. H. Cobb, *Proc. Nat. Acad. Sci. USA* **2018**, *115*, 3840-3845.
